# Non-catastrophic collapse of VEEV by lethal mutagenesis

**DOI:** 10.1101/2022.06.21.496950

**Authors:** Brian Alejandro, Eun Jung Kim, Jae Yeon Hwang, Juw Won Park, Melissa Smith, Donghoon Chung

**Affiliations:** Center for Predictive Medicine, University of Louisville, KY, USA; Department of Microbiology and Immunology, School of Medicine, University of Louisville, KY, USA; Department of Computer Science and Engineering, University of Louisville, KY, USA; KY INBRE Bioinformatics Core, University of Louisville, KY, USA; Department of Biochemistry and Molecular Genetics, School of Medicine, University of Louisville, KY, USA

## Abstract

RNA viruses replicate at a high mutation rate, providing a highly adaptive capacity to the population as well as vulnerability to an extinction due to additional mutations, which is the conceptual basis for lethal mutagenesis. However, the mechanistic understanding of lethal mutagenesis has remained unclear due to the lack of RNA mutagens with potent antiviral activity and the inherent inability to distinguish between infectious and non-infectious sub-populations within a viral population. In our study, we investigated how the replication competency and mutation frequency of Venezuelan Equine Encephalitis Virus (VEEV) change following treatment with the RNA mutagen β-d-N4-hydroxycytidine, a potent antiviral RNA mutagen. By pairing limiting dilution with a long-read sequencing approach, we specifically determined genomic sequences of replication-competent viral clones from the total population. We found that replication-competent VEEV population maintained itself within a narrow mutation spectrum with a significantly lower mutation frequency than the total population. We also found that treatment with an RNA mutagen did not induce catastrophic destruction of the infectious population even at a high exposure, allowing replication-competent subpopulations with increased variability in mutational fitness and enhanced adaptability to new selection pressures, such as antiviral treatment, to continue to propagate. Together our study provides new understanding of viral population landscape and suggests a careful consideration of lethal mutagenesis as an antiviral strategy for high-capacity replicating viruses such as alphaviruses.

## INTRODUCTION

Lethal mutagenesis is a process by which a population is pushed to extinction due to an increasing number of mutations(Bull et al., 2007). Since RNA viruses inherently replicate with a high mutation rate, a well-received antiviral approach has been to further increase the number of mutations by treatment with an RNA mutagen, especially for emerging RNA viruses that often lack specific treatment options. Historically, several antivirals, whose antiviral mechanisms were to induce lethal mutagenesis, have been developed, including ribavirin(D. H. Chung et al., 2007, 2013; Crotty et al., 2001; Jordan et al., 2000) and T-705 (a.k.a Favipiravir) (Baranovich et al., 2013; Furuta et al., 2017). They are known to exhibit their antiviral effect by increasing the mutation frequency via G substitution. These molecules, however, display a relatively low potency, narrow antiviral spectrum, or high degree of off-target effects; therefore, the realization of lethal mutagenesis as an effective antiviral approach had remained underdeveloped.

Compared to ribavirin or T-705, the recently developed β-d-N4-hydroxycytidine (NHC, initial metabolite of molnupiravir, Fig. 1) showed potent antiviral activity against a broad-spectrum of RNA viruses(Malone & Campbell, 2021; Urakova et al., 2017). NHC is converted to its active form (NHC-TP), and incorporated into RNA by the RNA-dependent RNA polymerase (RdRP) without causing stalling(Kabinger et al., 2021). NHC-MP incorporated into viral RNA induces transition mutations (preferentially C-to-U) in the nascent RNA strand by tautomerization. Lethal mutagenesis as an effective antiviral approach is exemplified by molnupiravir, which induces mutations in viral RNA and has been recently approved for SARS-CoV-2 treatment by the Food and Drug Administration (FDA). This FDA approval highlights the potential of lethal mutagenesis as an effective antiviral approach for other viruses. The potent antiviral activity and seemingly few off-target effects position NHC as an ideal molecule to study the mechanism and potential risks of lethal mutagenesis.

**Figure 1:**
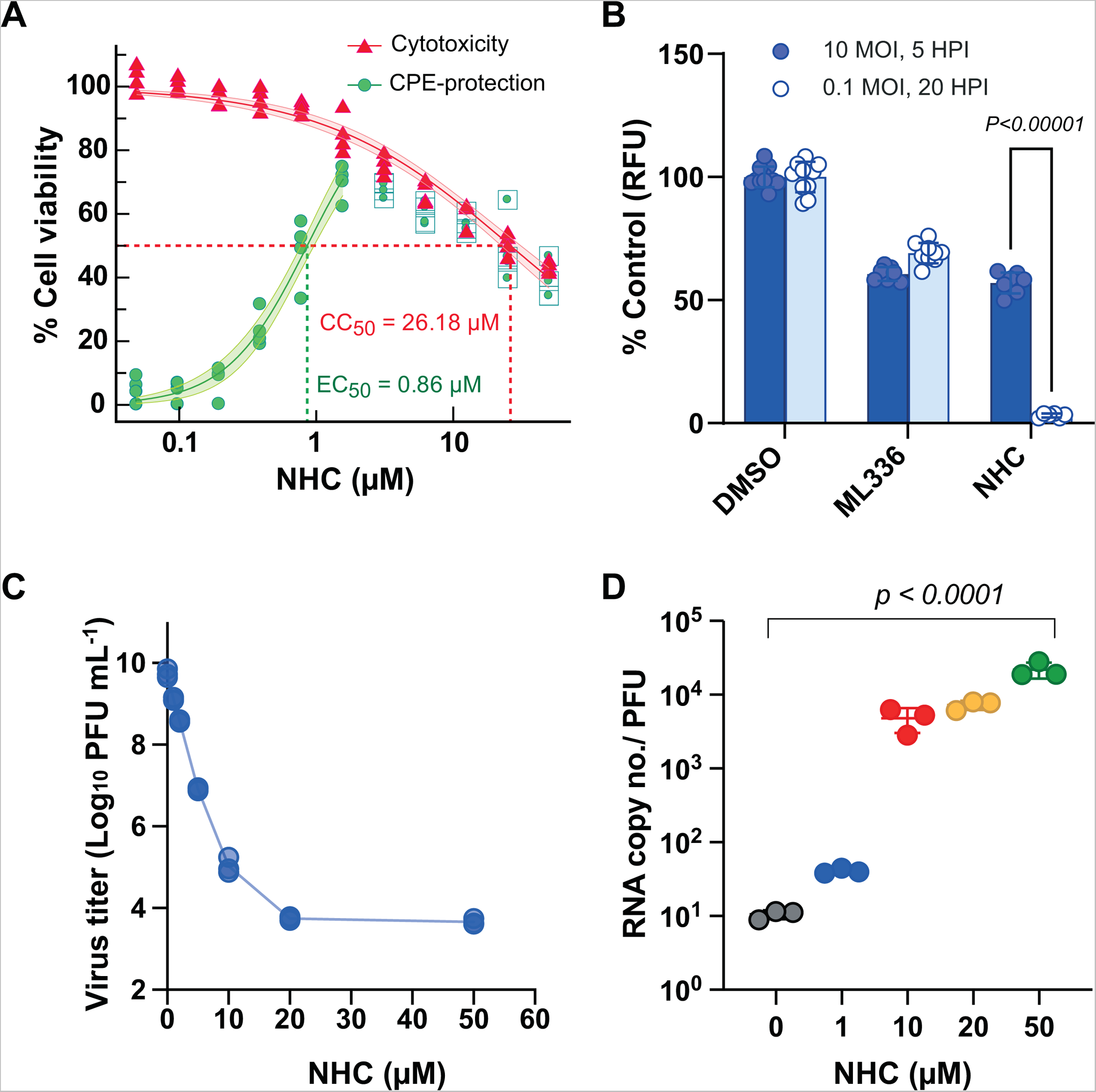
(A) Antiviral effect against VEEV TC-83 and cytotoxicity of NHC Normalized cell viability measured with Cell TiterGlo was depicted in red triangles for cytotoxicity(no virus infection) and in green circles for CPE-protection effect (with TC-83 infection) in Vero 76 cells. Values excluded from EC50 calculation were depicted with small green circles surrounded with a rectangle. The 95% confidence bands were shown as a shaded area. (n=4 per concentration) (B) Vero 76 cells grown in 96-well plates were infected with rVEEV TC-83-dR at two MOIs (10 MOI for a single cycle and 0.1 MOI for multi cycle replication) and treated with 50 nM of ML336 or 10 µM of NHC. The expression of EGFP signal was measured at the noted timepoints and normalized to the mean of the DMSO control (0.25%). Each dot represents independent wells. P value from Student t-test with n = 6). (C) Virus yield reduction by NHC. Virus titers of TC-83 cultured in the presence of NHC at the denoted concentrations for 18 hours were determined (n=3). (D) RNA copy numbers were determined by the realtime PCR method specific to nsP3 region with copy number standards. P< 0.0001, one-way ANOVA test.

Previous studies(Baranovich et al., 2013; Furuta et al., 2017; Sheahan et al., 2020; Yoon et al., 2018; Zhou et al., 2021)have clearly demonstrated that RNA mutagens induce an increase in mutations in viral genomes and seemingly display antiviral effects; however, thus far, no clear mechanistic demonstration as to how lethal mutagenesis actually leads to antiviral effects has been presented. In other words, a cause-and-effect relationship for lethal mutagenesis has not been clearly shown. Although it is well-accepted that RNA viruses replicate with a high mutation rate and mutations are generally disadvantageous for viral populations, several key questions especially regarding the infectious population (i.e., a group of replication-competent individual infectious units) before and after RNA mutagen treatment remain: 1) what are the nucleotide-level differences in viral genomes between infectious and non-infectious populations; 2) what is the mutation frequency and extent of heterogeneity in the normal infectious - not the total - virus population; 3) how much of an increase in mutation frequency is needed to achieve lethal mutagenesis; and 4) what is the minimum concentration of RNA mutagen needed to achieve the desired mutation frequency within the infectious populations? Addressing these questions will be critical for the continued development of more effective antivirals that act via lethal mutagenesis.

Previous studies have analyzed the genetic sequences of a total viral population treated with lethal mutagens without distinguishing between infectious and non-infectious sub-populations. Conventionally, mutation frequency (the number of detected alternate sequences/ total number of sequences analyzed) of a viral population is analyzed collectively by bulk sequencing of the total (replication-competent as well as incompetent) target population with a short-fragment next generation sequencing. This approach cannot identify or define the specific genomic diversity of the infectious populations due to bias towards sequences from the fittest or, conversely, non-viable populations. In addition, due to the use of short read sequencing methods, a comprehensive, quantitative mutational analysis for each genome or frameshift mutations is extremely limited by these methods.

To understand how lethal mutagenesis by RNA mutagen treatment leads to antiviral effects for RNA viruses, we sought to develop a new approach that focuses on characterizing the infectious portion of virus population. We model this approach using the Venezuelan equine encephalitis virus (VEEV) TC-83 strain, a member of the family Togaviridae. VEEV has an ~ 11 kb positive sense RNA genome and replicates at high titers in multiple cell lines. Various stable reporter viruses have been developed by others (Sun et al., 2014), allowing comprehensive examination of viral replication and genetic diversity.

In our study, we observed that infectious, replication-competent populations do demonstrate different profiles from the total population with respect to mutation frequency and response to RNA mutagen. We also found that catastrophic destruction of the infectious population did not occur even at a high exposure, leaving subpopulations infectious with increased variability that can lead to enhanced adaptation to new environments after RNA mutagen treatment.

## RESULTS

### Antiviral effect of NHC against VEEV TC-83

To study lethal mutagenesis, we first tested the antiviral effect of two well-known RNA mutagens, ribavirin and NHC, against VEEV TC-83 in a CPE-based assay. The antiviral effect of ribavirin was minimally detected as described by others(Markland et al., 2000) (data not shown); however, NHC showed a dose-dependent antiviral response in our assay with an EC_50_ of 0.86 µM and CC_50_ of 26.2 µM (**Fig. 1A**), which was consistent with data published by others(Urakova et al., 2017).

As NHC is an RNA mutagen, we further sought to understand how induction of mutations by NHC results in replication incompetent progeny virus, ultimately leading to detailing the NHC-mediated antiviral effect. We tested the antiviral effect of NHC using rVEEV TC-83-dR at two MOIs, MOI=10 for single round of replication and MOI=0.1 for multiple rounds of replication(Sun et al., 2014). This virus expressed EGFP in concert with the expression of the structural genes under a subgenomic promoter during replication. We found that while 10 µM of NHC showed only a 43.0% reduction in reporter signal compared to the control in the high MOI for assessing the single round replication impact, the antiviral effect for the second round of replication seen in the low MOI condition was significantly higher (97.9% reduction, n=6, **Fig. 1B**). This phenomenon was not detected with ML336, a direct antiviral against VEEV viral RNA synthesis(Skidmore et al., 2019). This result implies that NHC exhibits its antiviral activity not during viral replication but on progeny virus in subsequent rounds of replication, which is consistent with the concept of lethal mutagenesis.

### Mechanism of lethal mutagenesis by NHC

To understand how lethal mutagenesis works for VEEV TC-83, we investigated the effect of NHC during a single round of viral replication. We measured its antiviral activity in a virus yield reduction assay within a wide range of concentrations of NHC, from 1 to 50 µM, with TC-83 virus in Vero 76 cells (MOI ~1). The progeny virus titers were analyzed as a function of the concentration of the compound. Overall, NHC showed a significant reduction in virus titer compared to the mock-treated group (**Fig. 1C**) as previously reported by others(Urakova et al., 2017). However, interestingly, we found that the antiviral effect did not further increase at concentrations higher than 20 µM, resulting in a similar amount of replication-competent progeny virus titer at 50 µM. In other words, the antiviral effect of NHC was saturated at 20 µM and some fractions of the populations were still able to maintain their infectivity even with > 20 µM NHC treatment. However, such a saturation effect was not detected when we determined the relative infectivity (RNA copy number / pfu). RNA copy/pfu ratio of virus produced in the presence of NHC increased continuously up to 50 µM.

This result indicates that the overall quality of viral genomic RNA decreases due to NHC in a concentration-dependent manner; however, a significant fraction of virus remains replication-competent even at higher concentrations of NHC after a single round of replication.

### The replication-competent VEEV population displays a significantly lower mutation frequency than the total population

Our finding of replication-competent virus in the virus populations produced in the presence of NHC (NHC-conditioned population, hereafter) raised the question of how the error catastrophe model, in which a population collapses beyond its threshold of mutation rate, works and how increased mutation frequency affects viral infectivity(Eigen, 2002). To gain insight into the genetic structure of virus populations after RNA mutagen treatment, we developed two long-read based sequencing approaches for determining the genomic sequences of the infectious (Single Infectious Unit or SIU sequencing hereafter) and the total virus populations. The SIU approach is based on limiting dilution and multiplexed barcoded amplicon sequencing (**Fig. 2A**). Unlike conventional bulk sequencing that analyzes an entire virus population with multiple short-read sequences aligned to a template, our SIU approach only determines viral sequences of virus populations that can replicate at the individual infectious unit level (*Supportive Figure 1*).

**Figure 2:**
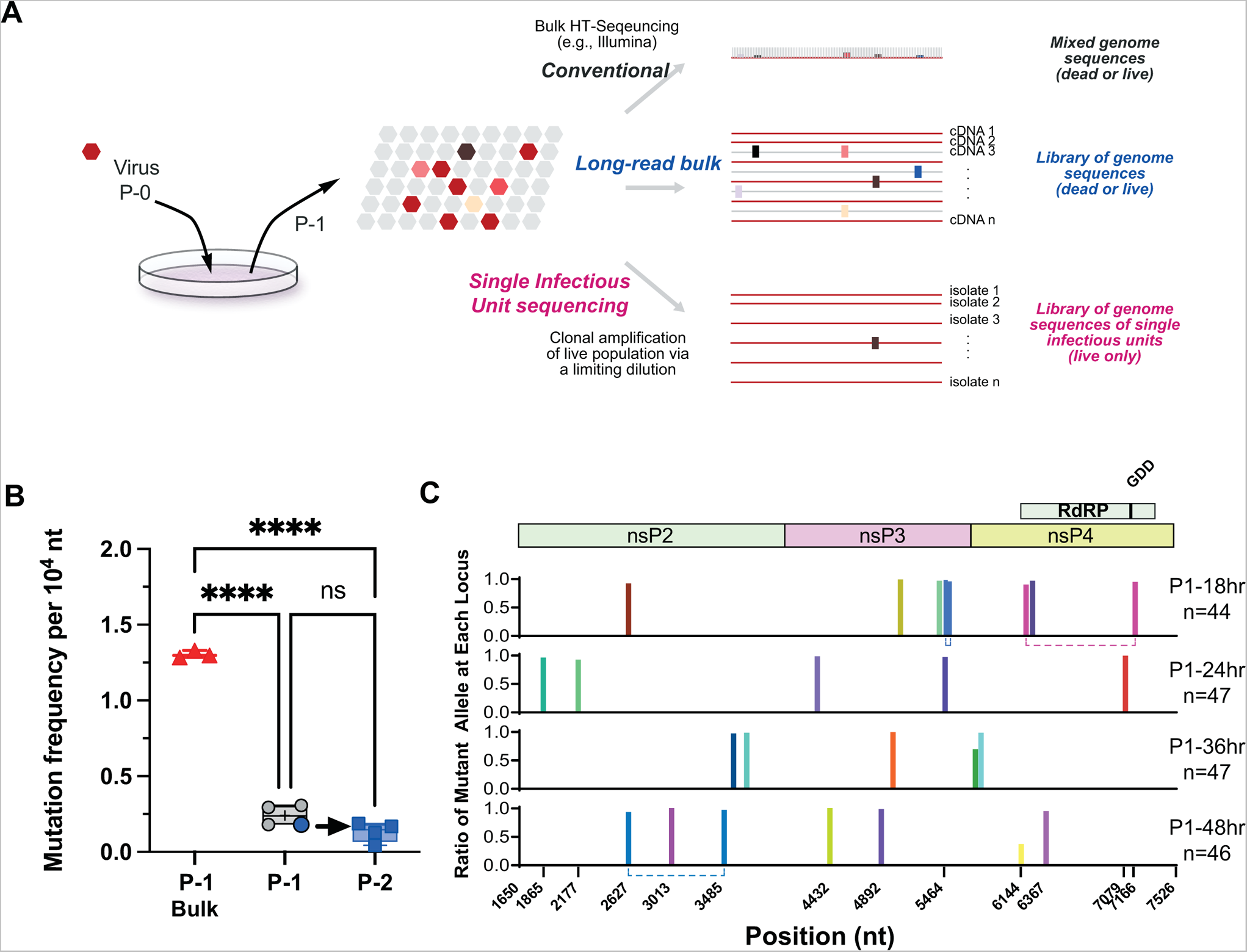
(A) Three different approaches for determining the genetic sequences of virus populations and their outcomes. (B) Comparison of mutation frequency between the total and infectious population with the bulk and SIU approach. The blue dot indicates the virus passed to the next round of replications **** p<0.0001, from one-way ANOVA test with Dunnett’s multiple comparison test. (C), Distribution mutations identified from the normal single infectious population of VEEV TC-83. Each color represents a single infectious unit isolated by a limiting dilution. Dotted lines indicate multiple mutations simultaneously and directly identified in a single infectious unit.

For total population characterization, long-read bulk sequencing was performed without the limiting dilution and selection of replication-competent clones; therefore, this approach generates collections of genomic sequences at a single molecule level regardless of their replication ability.

Using this strategy, we compared the mutation frequency of the replication-competent population of our VEEV TC-83 stock from that of the total population. For SIU sequencing, we tested 4 different cultures harvested at 18-, 24-, 36-, and 48-hours post infection to see if the mutation frequency of a virus population changes over its replication cycle (44-47 isolates per culture sample).

An overall analysis showed a much lower mutation frequency for the infectious population compared to the total population. The average mutation frequencies obtained from the SIU sequencing approach, 0.18 - 0.3 × 10^−4^ substitution per nucleotides (s/n), were approximately 10-fold lower than that from the bulk sequencing of the total population (1.30 × 10^−4^ s/n, n=3 **Fig. 2 B**), which is consistent with other viruses using similar approaches (e.g., sequencing of viral RNA produced from infected cells) for other RNA viruses (D. H. Chung et al., 2007; Drake, 1999; Jordan et al., 2000). This result clearly indicates that the replication-competent (i.e., infectious) fraction of virus population has a much lower mutation frequency than previously observed and it maintains a much less diverse population compared to the entire population. Assuming one round of infection (i.e., MOI > 1, and multiple harvests without a change in mutation frequency), our mutation frequency represents the mutation rate per generation.

To see if the long passage history of the stock virus we used affected mutation frequency and to measure the mutation rate per generation, we selected four single clones of virus from the clonal isolation procedure (P-1) and repeated the experiment. Two clones with no mutations, one with a synonymous mutation at nsP4 C459C, and one with a non-synonymous mutation at nsP3 E287D, were chosen for the experiment. This analysis showed that the frequency of mutants (2.6% −11.1%) and mutation frequency (0.04 - 0.19 × 10^−4^ s/n) of the infectious population were approximately 50% lower than those of P-1 samples; however, the differences were not statistically significant (**Fig. 2B**, and *Supportive Table 1*), indicating that the mutation frequency of infectious population is not significantly affected by a long history of continuous passage.

The SIU analysis showed that most of the infectious population (85 - 89%) had no mutations within the amplicon and only 10.6 - 15.2% of total isolates had one or more mutations different from the consensus sequence (*Supportive Table 1*). All mutants, except two, showed > 90% of reference nucleotide sequence at each location, confirming that they are indeed a single clonal isolate. No mutational hot spots were identified; the locations of mutations were randomly distributed throughout the amplicon except for the middle region (6367 - 7079 n.t.) of the RdRP domain in the nsP4, which did not accumulate any mutations (**Fig. 2C**). Since this is only a single passage, the mutation rate of TC-83 under normal condition is calculated to be 0.04 - 0.19 × 10^−4^ substitutions per nucleotide site per cell infection (s/n/c).

An analysis of the type of mutations within the replication-competent populations (***Table 1***) showed that non-synonymous mutations were approximately 2-fold more abundant than synonymous mutations (17 vs. 9) and transition mutations were found slightly more frequently than transversion mutations (14 vs. 12). Neither frameshift mutations, nor nonsense mutations were found in the viable populations.

Overall, our SIU sequencing approach indicated that the replication-competent population of VEEV TC-83 maintains its viability with a much lower mutation frequency (i.e., more homogeneous population) compared to the total population, and the majority of the infectious population did not have mutations within the nsP2-nsP4 region. This clearly shows that the diversity of RNA virus populations that can establish an infection is narrower than that previously determined with the total population.

### Genetic structure of replication-competent population of NHC-conditioned virus

Using our two approaches, we determined and compared mutations in the replication-competent and the total populations of viruses produced in the presence of NHC. The mutation frequency of the entire population determined by long-read bulk sequencing showed a significant increase in mutation frequency due to NHC treatment in a concentration-dependent manner. The mutation frequency of the total population increased to 32.65 × 10^−4^ s/n at 50 µM, which is approximately 25-fold increase compared to the DMSO control (1.30 × 10^−4^ s/n, n=3, **Fig. 3A**). For the replication-competent population of TC-83 cultured in NHC, however, the mutation frequency reached only to approximately 6.3 × 10^−4^ s/n, which is significantly lower than the total population (6.36 vs. 32.65 × 10^−4^, p<0.001, Two-way ANOVA test). Interestingly, the increase of mutation frequency was not concentration-dependent, reaching its maximum at 10 µM of NHC, without a significant increase at treatment concentrations higher than 10 µM.

**Figure 3.**
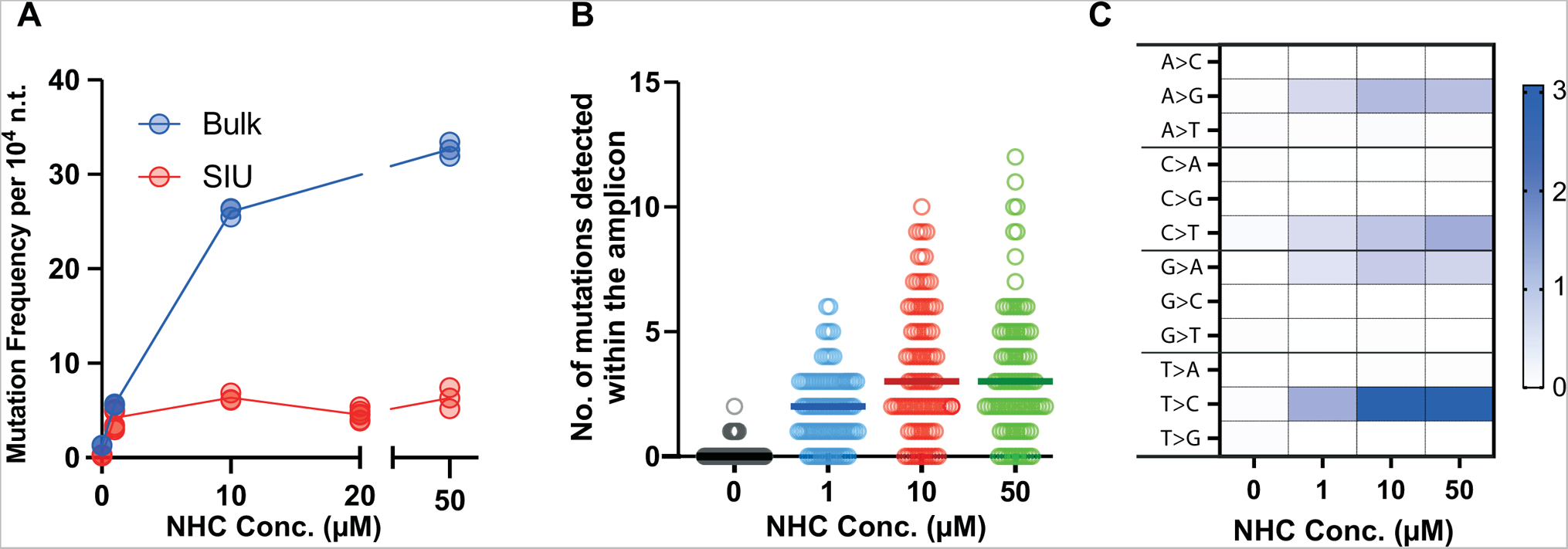
A) Mutation frequency of the total (blue) and infectious (red) progeny virus of VEEV TC-83 treated with NHC in Vero 76 cells. Each circle represents one biological sample (n = 3 ~ 6). B) Number of mutations detected in nsP2-nsP4 amplicons (5.8 kb) of progeny viruses. Replication competent progeny virus harvested from NHC treated cells was individually selected and its nsP2-nsP4 sequences were determined through the S.I.U. procedure. Each circle represents an individual replication-competent unit isolated after the limiting dilution and the thick lines represent the medians. C) Analysis of mutation types detected in the infectious population treated with NHC. Frequency of substitutions per 10,000 n.t. (n=3).

To understand more precisely how the mutations are distributed within the infectious population, we analyzed the number of mutations in each isolate (**Fig. 3B**, and **Table 1**). The median number of mutations from the 1 µM NHC-treated infectious population was 2 within the 5.9 kb amplicon examined, compared to 0 from the DMSO-treated group. While the median number of mutations increased only marginally (from 2 to 3) when treated with ≥ 10 µM of NHC, the change in the distribution of the number of mutations in the amplicon treated with NHC was more obvious, especially for the strands with more than 5 mutations within the amplicon (6.8% and ≥ 30% for 1 and 10 µM NHC-conditioned group, respectively)

**Table 1.**
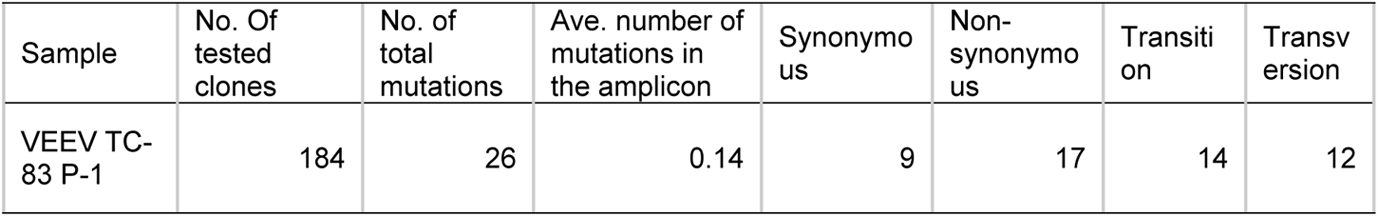
Analysis of mutations within the nsP2-nsP4 amplicon from the replication-competent population of VEEV TC-83

To our surprise, we found significant fractions of NHC-conditioned, replication-competent viruses with a low number of mutations. The frequencies of infectious units with mutations less than 2 within the amplicon comprised of 63.6%, 44.3%, and 45.5% of their populations in the 1,10, and 50 µM conditioned groups, respectively. Particularly, more than 12.5% of the replication-competent population was mutation-free in all NHC-conditioned groups (**Table 2**).

**Table 2.**
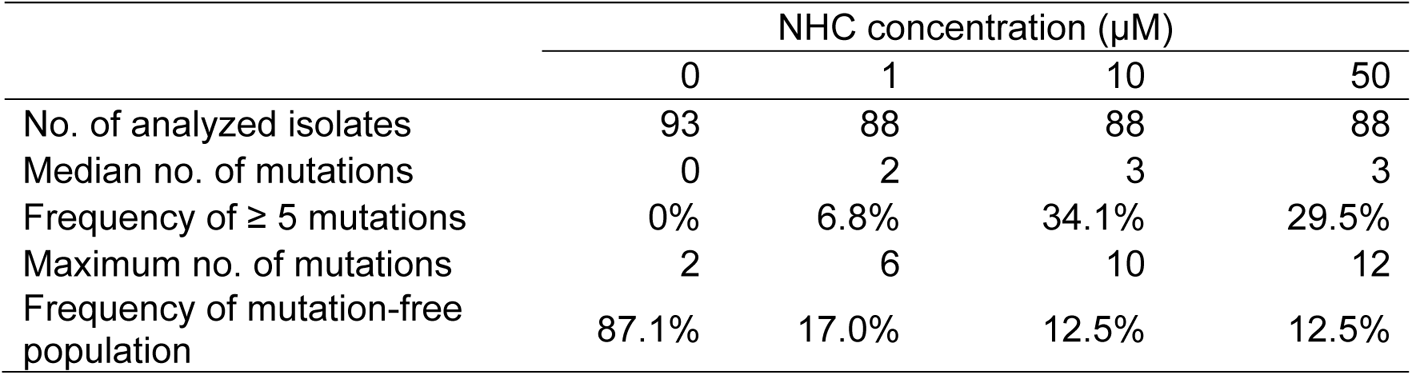
Distribution of mutations within the nsP2-nsP4 amplicon from the infectious population of VEEV TC-83 treated with NHC

An analysis of mutations identified in the infectious population (Fig 3.C) showed that all four types of mutations that can be induced by NHC(Sheahan et al., 2020), indicating the incorporation of NHC into viral RNA might have happened during both (-)-sense RNA and progeny genomic RNA synthesis. However, interestingly, the majority of mutations induced by NHC in VEEV were T-to-C (U-to-C in the viral genome), indicating the incorporation of the NHC occurred during the synthesis of genomic RNA across adenosine (***Fig. 3C***). This is a different finding from the SARS-CoV2 study, performed with the total RNA sequencing, showing a preference of G-to-A or C-to-T mutations by NHC(Sheahan et al., 2020).

Overall, our data showed that the effect of the NHC RNA mutagen on the mutation frequency of replication-competent viruses is not as drastic as that observed in the total population. Instead, the higher mutation rate condition induced by RNA mutagen treatment increased the diversity of the replication-competent population. Importantly, the population did not experience a catastrophic collapse during single cycle replication, leaving significant populations with a low number of mutations or mutation-free.

### NHC-treated, replication-competent virus shows a retarded yet a diverse phenotype in replication

Based on our finding that NHC-conditioned VEEV still have replication-competent population with increased diversity (i.e., mutated genome), we sought to measure the extent of growth defects induced by those mutations acquired by the population. To test this, we established a viral growth kinetic assay and analysis pipeline for single infectious units using rVEEV TC-83-dR. To evaluate growth variations, we determined the maximum growth rate (gr, the maximum slope of EGFP expression) and the Area Under the Curve (AUC) of the EGFP expression from individual wells infected with virus at 0.5 TCID50/well. Under this condition, approximately 20 – 30 percent of wells were positive for viral infection, implying that the majority of virus-positive wells were a single isolate. We found that indeed NHC-conditioned virus showed a significant decrease in viral replication with a decrease in gr and AUC with a negative skew in the histogram analysis (**Fig. 4 B and C**). For example, the growth rate and AUC of the NHC 10 µM-conditioned group was reduced by 20% compared to the mock group. Inversely, other time-associated kinetic parameters such as the lag time (the time intercept between the regression line associated with the gr calculation and the baseline defined by the first point in the calculation zone, defined as ‘LagC’), and time at maximum growth rate (t_gr) were increased, indicating a delayed replication of the NHC-conditioned populations compared to the mock-treated group (*Supportive Figure 2*). At the individual level, however, approximately 44 – 55 % of the single isolates from NHC-conditioned virus showed growth rates similar to the untreated group (mean ± 1 × S.D).

**Figure 4.**
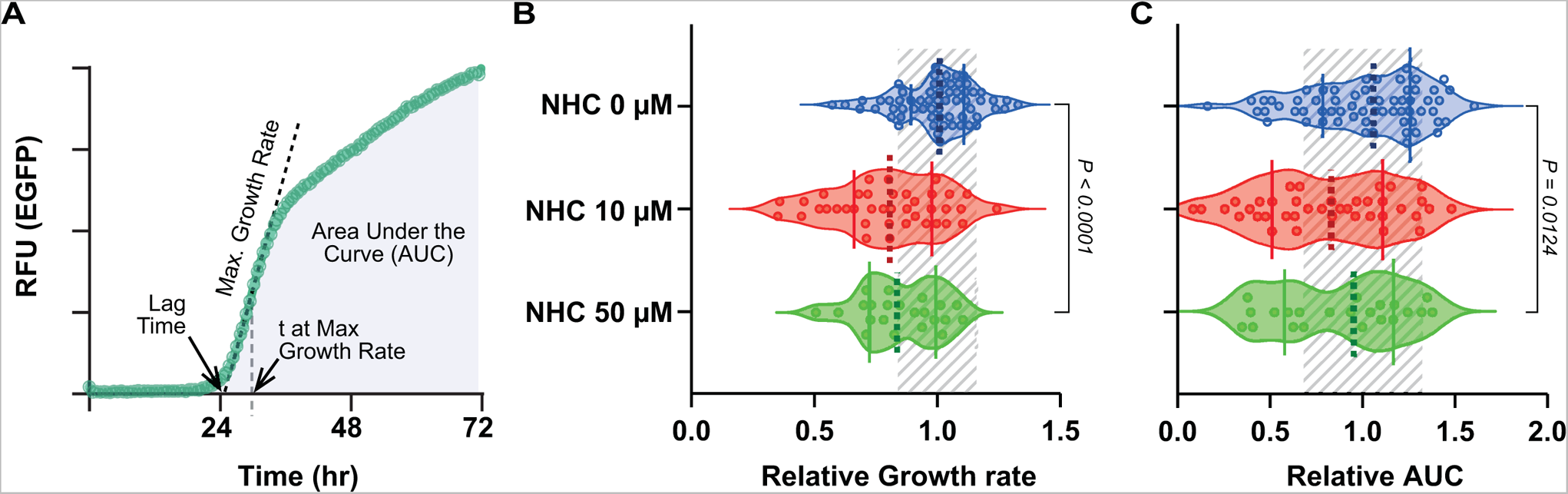
***A***) Single infectious unit growth analysis. Cells plated in a 384-well plate were infected with rVEEV TC-83-dR at 0.5 TCID50/well and the expression of EGFP was monitored in a plate reader with a live culture mode. (A) A representative growth curve of rVEEV TC-83-dR shown for demonstration of growth parameters. Distribution of growth rate (gr) (B) and AUC (C) of mock- or NHC-conditioned infectious populations. P values were from one-way ANOVA test with Brown-Forsythe method.) (n= 63,39, and 22 for the mock, 10, and 50 µM NHC treated groups. Dotted and solid lines represent the median and top and bottom 25% percentile in each group and the shaded areas indicate mean ± 1 × S.D from the mean of the mock-treated group.

These results demonstrate that the replication-competent virus population produced from treatment with NHC has a negatively skewed, but not universally shifted growth rate distribution.

### RNA Mutagens May Enhance Viral Adaptation to New Environments

So far, our data showed that an increase in mutation rate does not induce a catastrophic destruction of the population; rather, it leads to some replication-competent populations with varying, yet limited, degree of diversity. An increase in diversity within a virus population (i.e., population complexity) by an RNA mutagen may instead enhance adaptation of the virus population to a new environment under different selective pressures. Here, we sought to test this hypothesis in the context of adaptions that benefit their survival, particularly in therapeutic situations such as conferring resistance to other direct-acting antivirals. We employed ML336, which is a direct-acting alphavirus inhibitor targeting the nsP2 and viral polymerase nsP4, inhibiting viral RNA synthesis(D. Chung et al., 2012; D. H. Chung et al., 2014, p. 2; Schroeder et al., 2014; Skidmore et al., 2019, p. 2). ML336 has a relatively low resistance threshold, conferring a phenotypic resistance in EC_50_ greater than 100-fold with a single nucleotide change at key locations such as nsP2 Y102 or nsP4 Q210 without continuous passaging. These characteristics establish ML336 as an ideal marker molecule for investigating the impact of mutagenesis from RNA mutagen treatment on sensitivity to direct-acting antiviral compounds.

In a preliminary study with selection of ML336-resistant clones by treatment with 3 µM of ML336 ( ~ 100 × EC_50_) in agarose overlay, we found an approximately 10-fold increase in the frequency of ML336 resistant viruses derived from 1 or 10 µM NHC-conditioned virus stocks (2.67 vs. 24.6 per 10,000 pfu, data not shown).

To obtain statistically improved data with a scaled-up experiment, we utilized a limiting diluting method with a modification of the SIU approach where 100 and 5 pfu/well were used for the untreated and NHC-treated populations, respectively. Based on our preliminary agarose overlay experiment, the expected frequency of ML336 resistant isolates in this condition was < 0.002 per well, which is low enough to isolate a single resistant clone per well. Infected cells were cultured in the presence of 1 µM of ML336 (~ 30 × EC_50_) in the culture media and the number of wells that showed signs of infection were counted as ML336-resistant.

The mock-treated population showed ML336-resistant frequency of 15.27 ± 2.96 per 10,000 pfu (mean ± SEM, n= 7 from two independent experiments). Compared to this, the viruses produced in the presence of 1 and 10 µM of NHC showed a ML336 resistance frequency of 60.46 ± 5.92, and 193.2 ± 25.09 (mean ± SEM, n= 7 from two independent experiments), respectively, showing a strong dose-dependent increase in ML336 resistance by 4 and 12.6-fold compared to the non-treated group (*p < 0.005*, One way ANOVA test, see ***Fig. 5A***).

**Figure 5.**
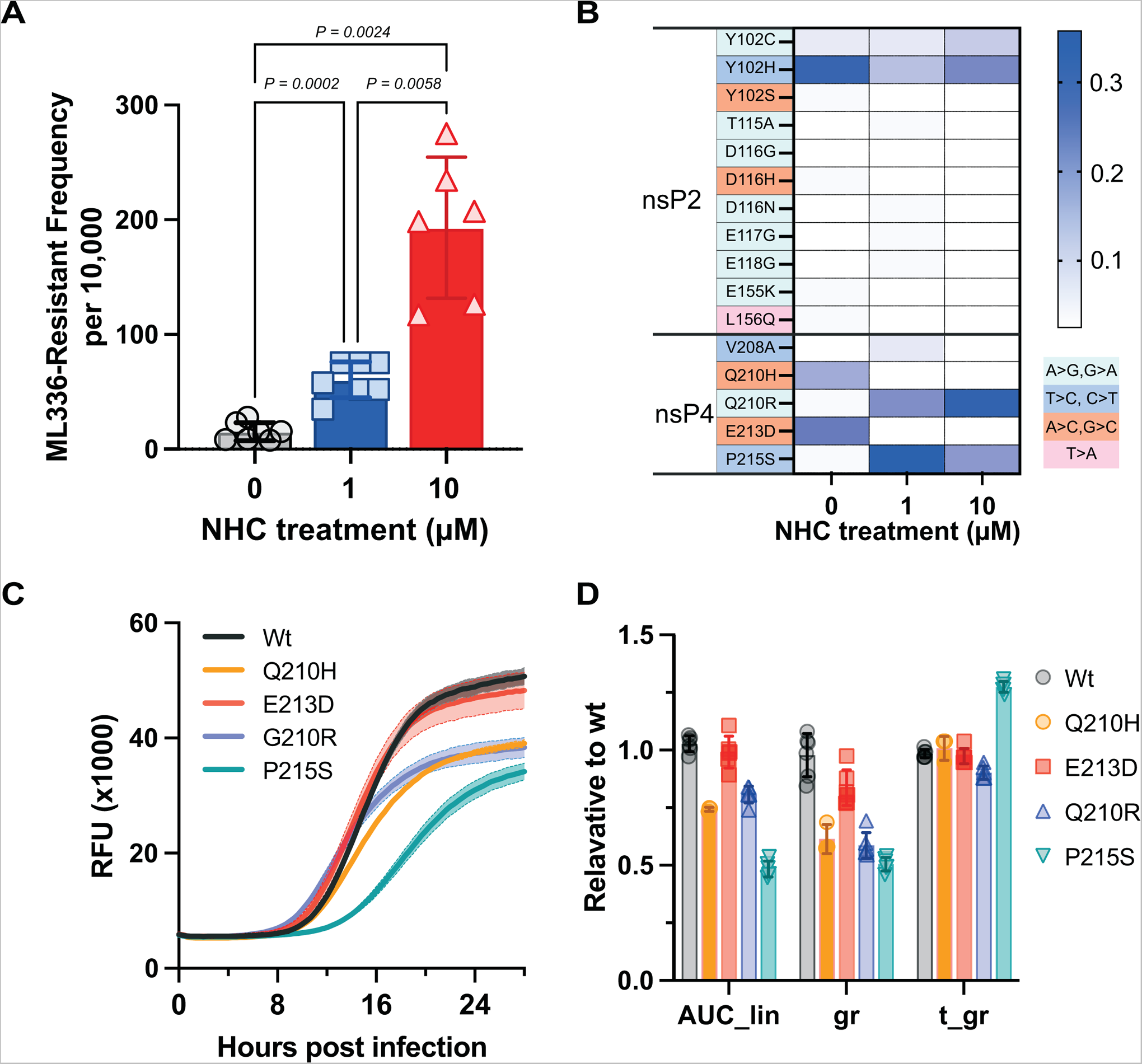
Increase of frequency of resistant virus against a direct acting antiviral by NHC. ***A)*** Frequency of ML336-resistant virus in NHC-conditioned VEEV TC-83. Progeny viruses cultured in the presence of NHC were screened for resistant mutants to ML336 (1 µM). P values are from one-way ANOVA test with Dunnett’s multiple comparison test. Means (bars) and standard deviations (error line) are shown with each data point. ***B***) Analysis of mutations found in ML336 resistant VEEV TC-83 strains generated in the presence of NHC. Frequency of mutants with designated mutations (n= 29,28 and 40 isolates for 0,1, and 10 µM NHC-conditioned VEEV TC-83). ***C)*** Growth curve analyses of mutant strains. Mutations were introduced to rVEEV TC-83-dR and the growth kinetics analysis was conducted at 0.1 MOI for 28 hours. Bold lines and shadowed area represent the means and the error enveloped derived from the standard deviations (n=6) except Q210H (n=3). ***D)*** AUC_lin : Area under the curve; gr: maximum growth rate; t_gr: time at the maximum growth rate.

Next, to confirm the increase in the frequency of ML336-resistant mutants was due to mutations induced by NHC, we compared the genomic sequences of ML336-resistant clones derived from the mock treatment to those from the NHC treatment group. We found a significant change in the population landscape of ML336 resistant group. (***Fig. 5B***). ML336-resistant clones isolated from the mock treatment group showed a preference to nsP2 Y102H (T>C, transition mutation, 31%), and nsP4 Q210H and E213D (A>C, transversion mutations, 17.2 and 24% respectively). In ML336-resistant viruses isolated from the NHC-conditioned viruses, however, two mutations that were not selected in the non NHC-conditioned group, nsP4 Q201R (A>G, transition mutation, 21%) and P215S (C>T, transition mutation, 38%), were isolated as a dominant variant seen. Few isolates with a transversion mutation were identified within the ML336-resistant isolates that had originated from NHC-conditioned groups. Notably, NHC drives the population with transition mutations, supporting the idea that increased diversity by NHC-induced transition mutations facilitated a different subset of the ML336 resistant mutants that were not evident in the normal population. To validate if the newly isolated mutations confer resistance to ML336, we evaluated phenotypic resistance of ML336-resistant mutations that were uniquely identified from NHC-conditioned population (**Table 3**). We found that nsP4 Q210R and P215S showed EC_50_s of >100 and 18.6 µM, demonstrating potent resistance to ML336 treatment.

**Table 3.**
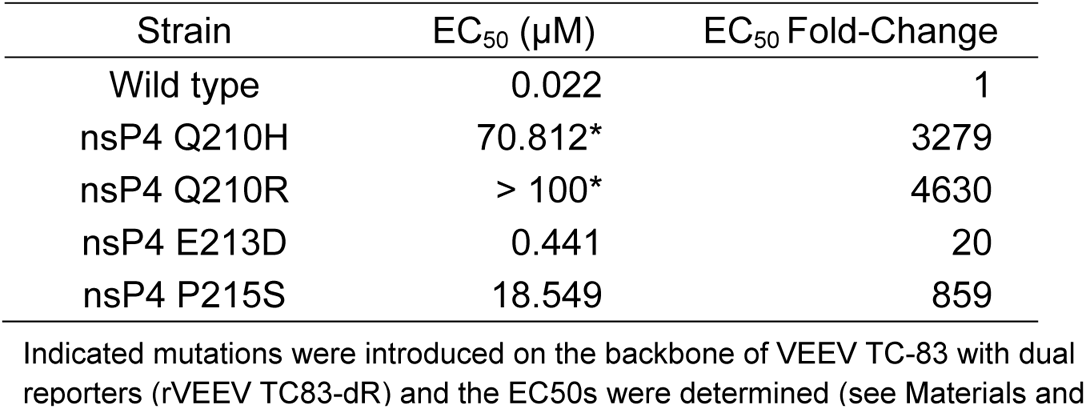
Phenotypic resistance of ML336-resistant mutants uniquely identified by the treatment of NHC.

To understand if the two ML336-resistant mutant isolates (nsP4 Q210R and P215S) have a lower fitness than others, resulting in a lower frequency in the untreated, ML336-resistant group, we analyzed the effects of the mutations on viral replication using our rVEEV TC-83-dR reporter system (***Fig. 5C and D***). nsP4 Q210H and E213D mutant clones displayed similar or attenuated growth profile compared to the wild type (74.3 and 99.1% in AUC_lin). However, nsP4 Q210R and P215S showed a bigger growth defect with 80.1 and 48.4% in AUC_lin compared to the wild type. This experiment showed that a diversified and/or skewed viral population from an increased mutation rate condition may provide an opportunity for less fit viral populations to emerge under certain pressures.

Overall, these experiments clearly showed that increased diversity of RNA virus populations may provide higher adaptation competency to new environments by expanding the genetic landscape.

## DISCUSSION

Our study shows how RNA mutagen affects the genetic and phenotypic landscape of the replication-competent subpopulation after treatment. We also show that the replication-competent VEEV TC-83 virus population has a narrow range of genetic diversity, and this limited population becomes diverse, yet much lower than the mutational frequency of the total viral population (replication-competent, and -incompetent combined), under artificial conditions that lead to an elevated mutation rate.

### Viral extinction by RNA mutagen treatment

Previous studies have demonstrated potent in vitro antiviral activity of NHC for multiple RNA viruses including VEEV (Agostini et al., 2019; Stuyver et al., 2003; Urakova et al., 2017; Yoon et al., 2018), which may initially appear contradictory to our finding of residual infectious virus after high concentration NHC treatment. Most of these studies focused on NHC’s antiviral activity in various viruses; hence, the antiviral activities were usually expressed as comparison to mock controls, or tested concentrations were within a lower range. Because our focus was to understand how RNA mutagens change exposed, progeny virus, we simply paid attention to the replication-competent population as a stand-alone experimental group, which is less than 10^6^-fold lower than the mock-treated group. Therefore, our results agree with previous findings of strong antiviral activity of NHC. In fact the previous publications showed non-complete inhibition of viral replication by NHC in vitro and in vivo across RNA viruses, except for coronaviruses(Sheahan et al., 2020; Zhou et al., 2021).

NHC showed a near complete inhibition of SARS-CoV2 and MERS in vitro and in animal models, demonstrating clearance of viral infection after NHC treatment at 10 µM(Sheahan et al., 2020), which is different from our results. This could be due to several reasons. Firstly, NHC may have a greater antiviral effect on coronaviruses, as their genome size is bigger. Although coronaviruses have a proofreading function encoded by the exoribonuclease domain (ExoN) in their non-structural protein(Smith et al., 2013), a longer genome size provides a greater opportunity for an RNA mutagen to get incorporated during replication, leading to overall decrease in viral fitness. Secondly, the size of infectious coronavirus pool is much smaller than that of the alphavirus. While typical virus titers of coronaviruses in cell cultures do not exceed 10^6^ pfu/mL, VEEV TC-83 replicates robustly and produces high viral titer progeny virus (> 10^8^ pfu/mL). Therefore, our model virus might have revealed an infectious population that would not be detected even with a similar level of antiviral activity. Thirdly, our study focused on a single round of replication to clarify mechanistic understanding, which may not reveal the consequential outcome from continuous treatment over multiple rounds of viral replication. Because accumulation of mutations negatively affects viral fitness overall, the full antiviral activity of RNA mutagens may need several rounds of viral replication to drive the entire population to become non-infectious.

### Difference between replication-competent and total population

Mutational frequency of RNA viruses have been measured largely by aligning sequences of small fragments of viral genomes harvested from infected cells or their supernatants. While the conventional approach is still appropriate to study mutation rates and other genetic features, the understanding of the genetic landscape of the replication-competent population, which contributes to the sustainability and fitness of viral infection, remains unknown for most RNA viruses. Here we developed a higher-throughput approach that is based on limiting dilution and long-read, multiplexed sequencing to quickly determine the consensus sequences of each replication-competent isolate. Our data showed that the mutation frequency of the VEEV replication-competent population (0.2 - 0.33 / genome or 0.18 - 0.3 × 10^−4^ s/n) is approximately 10-fold lower than that from the entire population or values previously determined by others. This finding implies that the virus population is maintained by constituents with a small number of mutations. In other words, progeny virus with a high number of mutations is not capable of replicating and this could be a reason for low specific infectivity (pfu/copy number of viral genome) in addition to other observations, such as the absence of RNA capping in the viral genome(LaPointe et al., 2018). The replication incompetence of the high-mutation-containing population might interfere with the replication-competent members by competing for viral receptors or inducing innate immune responses.

### Mutation frequency threshold at the population level

RNA virus populations are believed to replicate on the edge of error catastrophe, where the virus population may experience an extinction event due to increased mutation rate. This hypothesis served as the basis for the theory of lethal mutagenesis(Crotty et al., 2001; Eigen, 2002; Malone & Campbell, 2021). However, no thresholds at which virus populations could preserve their infectivity have been clearly defined. This could be due to the lack of effective RNA mutagens or the low resolution of the methods that have been used to determine mutational landscapes in the replication-competent populations. In our study, we addressed this with NHC, an effective RNA mutagen with antiviral activity against a broad-spectrum of RNA viruses, as a tool to modulate the viral error rate in the system. In our system, we did not observe a catastrophic collapse of the population even at the maximum treatment concentration (50 µM). Instead, we found populations that were able to replicate with a high mutation rate. Interestingly, the mutation frequency of these viruses did not surpass 6.3 × 10^−4^ s/n, roughly 6.5 mutations per genome at 10 µM of NHC treatment. At the population level, our data implies that even under a high mutation condition, a small fraction of virus remains productive while the majority of total progeny viruses are no longer productive due to accumulated mutations that occur in a concentration-dependent manner. This is seemingly different from the lethal mutagenesis theory; however, this difference might be due to the design of study using a single round infection and the high population size of VEEV (> 1×10^8^ pfu/mL).

### Mutation frequency at the individual level

Our system can provide a high resolution mutational landscape of the population at the individual RNA or infectious unit level by taking advantage of long-read sequencing approaches(Amarasinghe et al., 2020; Burgess, 2018; Pollard et al., 2018). Unlike short-read based methods (e.g., Illumina sequencing platform), long-read sequencing can provide the precise number of mutations for each strand in phase. Therefore, even though sequencing read depth may be lower than with short-read approaches, the increased resolution and low error rates without bioinformatic imputation or reconstruction of genomes is highly valuable. In our study, we determined the progeny viral genomic RNA sequences for the nsP2-nsP4 region of VEEV after replication in the presence of NHC. We demonstrated that NHC further diversified the replication-competent population. At the single replication-competent unit level, the power of our approach was highlighted by identification of a replication-competent clone with 12 unique mutations and characterization that 12.5% of the replication-competent population showed no mutations within the amplicon (approximately 50% of the total genome, Table 1). These types of analyses cannot be achieved by short-read approaches, and we believe our data provides a sequence landscape reflective of true viral population diversity reality.

### Mutational fitness effects

Overall, our comparative analysis of total vs. replication-competent populations implied a negative, deleterious effect of mutations, and agrees with previous notions of mutational fitness effects (MFE). However, our work further demonstrates the MFE at the level of individual viruses that were generated with increased mutation rates. In general, mutations are perceived to negatively affect viral replications. Pilar Domingo-Calap et al. analyzed the mutational burden and effect on viral survival from viruses using a random in vitro mutagenesis approach, and showed an overall deleterious, rather than beneficial, effect of mutations on viral fitness(Domingo-Calap et al., 2009; Sanjuán, 2010). However, studies of MFE on natural viral isolates are limited. For VEEV, a low fidelity mutant virus strain displayed insignificant changes in viral fitness and reduced virulence(Kautz et al., 2018). In the case of poliovirus, increased replication fidelity has shown to reduce both fitness and virulence especially in an animal model(Pfeiffer & Kirkegaard, 2003, 2005). In our study we sought to determine the precise relationship between mutations and growth profile when the virus is produced with a higher mutation rate in the presence of NHC. It should be noted that the majority of virus produced from NHC treatment were replication-incompetent as shown in the titer reduction experiment (Fig 2C). However, there remained some proportion of replication competent progeny virus. Our approach to determine the effect on the fitness was to utilize EGFP expression as a surrogate marker of viral structural protein expression in infected cells (Sun et al., 2014)and analyze the growth profile of viruses derived from NHC-treated populations. EGFP signal is a cost-effective readout and can be continuously monitored in real-time in a high-throughput manner. Considering that the half-life of EGFP has been reported as approximately 83 min (< 1.5 hrs). in cells(Danhier et al., 2015), the EGFP signal we measured is considered to have originated from the EGFP accumulated between the time of reading and 3 hours prior. Therefore, EGFP can act as a reasonable readout for viral replication in this system. We have investigated various kinetic parameters that could differentiate growth patterns best using PCA analysis with all identified single clones identified in the study and found that normalized AUC, gr, lagC, and t-gr are the biggest contributors to the principal components (data not shown).

NHC-conditioned, replication-competent virus groups displayed viral growth defects with an increase in lagC and a decrease in gr and AUC at the group level (Fig 4. B, and C). Considering the mutation frequencies of these groups were 6.3 × 10^−4^ s/n or ~ 20-fold higher than the non-treated group (0.18 - 0.3 × 10^−4^ s/n), the growth defects were considered less prominent than expected. More interestingly, a significant number of viral isolates within the NHC-conditioned group displayed a growth profile equivalent to the mock treated group (Fig4 B and C) and this finding agrees with our mutational analyses, showing ≥ 30% of population free of mutations within the nsP2-4 amplicon (Fig 3. B). This result could be because our analysis is selective for replication-competent isolates.

Overall these findings imply that 1) replication-competent viruses can still be selected from high mutation rate conditions, 2) virus isolates with multiple mutations can be replication-competent, 3) viral clones within a population may not be universally affected by mutagen treatment, but rather show a skewed frequency spectrum, and 4) mutation-free populations are present even in a higher mutation rate environments and can effectively serve as founder isolates of the next round of replication.

### Biological effects of increased diversity

Viral adaptation to new environments is a process of genetic fixation of beneficial mutations. The high mutation rate of RNA viruses is considered a primary reason for the highly adaptable characteristics of RNA viral genomes, as this high mutation rate provides a diverse population that can be selected from. From this perspective, an increase in population diversity may contribute to an increase in viral adaptation to new environments such as antiviral treatments or neutralizing antibodies. In our study we employed ML336, a direct acting anti-VEEV antiviral, to provide low threshold selection pressure and showed that an increase in population diversity indeed led to a higher incidence of resistant clones by up to 12-fold (Fig 5). The ML336-resistant clones derived from NHC-conditioned groups had mutational profiles different from naturally occurring ML336 resistant variants. This finding implies that treatment with RNA mutagens may shift the progeny population into a more diverse and transition mutation-rich population, leading to the emergence of novel isolates.

The nsP4 Q210R and P215S mutations showed a similar phenotypic resistance to ML336; however, these were not selected from the normal population, and uniquely identified under NHC-conditioned groups. This could be explained by the fact that these resistant populations showed an attenuated growth profile compared to the others, as we showed in Fig 5. A follow-up study will be needed to investigate if the novel resistant clones identified here become fixed in the outgrowth population or become dominant when they replicate among other isolates.

Together our study showed that the replication-competent VEEV RNA virus population maintained itself within a narrow mutation spectrum, and treatment with RNA mutagen resulted in two subpopulations: 1) replication-incompetent majority, and 2) a diversified, yet replication-competent, minor population, suggesting a careful consideration of lethal mutagenesis as an antiviral strategy for high-capacity replicating viruses such as alphaviruses.

## METHOD AND MATERIALS

### Cell culture and viral strains

Baby hamster kidney (BHK) clone 21 cells (ATCC CCL-10) and Vero 76 (ATCC® CRL-1587™) were maintained in Modified Eagle’s Medium with Earle’s Balanced Salt Solution and L-glutamine (MEM-E) supplemented with 10% fetal bovine serum (FBS) (Corning CellGro). Cells were maintained at 37 °C in humidified incubators with 5% CO_2._ VEEV TC-83 was derived from a lyophilized vaccine stock (USAMRIID, gift from Dr. Connie Schmaljohn). Reconstituted virus underwent a single round of replication in BHK C21, prior to storage at −80 °C until use.

VEEV TC-83 with dual reporters (rVEEV TC83-dR) were constructed with a plasmid backbone of pTC83-eGFP (gift from Dr. Kevin Sokoloski). In this construct, the entire cDNA of the TC-83(Sun et al., 2014) genome is under an SP6 promoter and eGFP is controlled under the double subgenomic promoter. The nanoluciferase gene was inserted at FseI site using the Gibson cloning method (pTC83-dR). Specific mutant viruses were generated using Quick-change mutagenesis with the pTC83-dR template (See *Supportive Method 1* for the primer sequences). The full sequence of the resultant clone was validated prior to rescue virus. Virus rescue was performed as described previously(Bernard et al., 2000).

### Antiviral assays

Anti-VEEV activity of NHC was measured using a cell-based CPE assay as previously described (D.-H. Chung et al., 2016). Compounds were dissolved in DMSO at 20 mM and stored at −20 °C. Briefly, Vero 76 cells seeded in a 96 well plate one day before were infected with virus at an MOI of 0.05 in the presence of test compounds serially diluted to 8 different concentrations. The final concentration of DMSO was maintained at 0.25%. Infected cells were incubated for 48 hours (hrs) prior to measurement of cell viability protected from VEEV-induced CPE was performed using CellTiter-Glo (Promega).

For phenotypic resistance, the EC_50_s of ML336, an anti-VEEV direct acting antiviral compound inhibiting viral RNA synthesis(Skidmore et al., 2019), against various strains were determined by using EGFP fluorescence signal (RFU) expressed from infected cells as a surrogate readout of virus replication. Vero 76 cells plated on solid black 96-well plates were treated with serially diluted ML336 as above and then infected with mutant rVEEV TC83-dR strains at an MOI of 0.05. After 24 hrs incubation at 37 °C with 5% CO_2_, cell culture media was replaced with 70 µL of PBS, followed by EGFP signal detection measured with the Synergy 4 (BioTek). EC_50_s were calculated with a 4-parameter logistic model using XLfit software (IBDS).

### Generation of NHC-conditioned VEEV TC-83

Vero 76 cells grown in T-25 flasks or 6-well plates were infected by incubating the cells with TC-83 at MOI of 1 for one hour at 37 °C. The cells were washed with PBS twice and replenished with virus growth media containing NHC. After 24 hours of incubation, the cell supernatant was collected and centrifuged to remove the cell debris. Virus was aliquoted and stored at −80 °C until use.

### Clonal isolation by limiting dilution

Vero cells were seeded in 384-wells at a density of 4000 cells/well in a volume of 15 µL and cultured overnight. Cells were infected with 0.1 pfu/well for normal population studies. Three days later wells with productive infection were identified by evaluating CPE using alamarBlue (BioRad). Cell supernatants of wells in which alamarBlue fluorescence signal was less than 4×standard deviation (sdev) of signal from uninfected cell controls were consolidated with a unique ID and then the virus clones were amplified once by infecting and culturing in 96-well plates seeded with fresh Vero 76 cells for 18 hrs.

### Bulk sequencing

RNA was isolated from infected cell supernatants with DirectRNAZol MagBead RNA (Zymo research). RNA was reverse transcribed to cDNA with Maxima reverse transcriptase primed using random hexamers and following the manufacturer’s protocol.

Total RNA isolated from 0.15 mL progeny virus was subjected to cDNA synthesis followed by targeted amplification of a fragment containing nsP2-nsP3-nsP4 (5.9 kb) with Phusion High-Fidelity DNA polymerase and uniquely barcoded primers. For multiplexing of samples, 9nt barcodes were used. The barcode sequences were generated by using an R package, DNA barcode (https://bioconductor.org/packages/release/bioc/vignettes/DNABarcodes/inst/doc/DNABarcodes.html) using an argument of mySeqlevSet <- create.dnabarcodes(9, metric=“seqlev”, heuristic=“ashlock”, cores=12). A total of 612 barcodes sequences were obtained and unique sequences were added to the 5′ end of the sequences for nsP2-fwd and nsP4 rev. Barcoded primers were purchased from IDT DNA. The primer sequences are listed in Supplementary data (Sequences of the barcoded_primers.xlsx).

PCR amplicons were purified with KAPA Pure magnetic beads (Roche) at a 0.5 × bead-to-sample volume ratio and eluted in nuclease-free water. Pooled barcoded PCR amplicons were prepared into SMRTbell template libraries as recommended by the manufacturer and subjected to single molecule, real time (SMRT) sequencing on the Sequel IIe system (Pacific Biosciences) using v2.0 chemistry and 30 hrs movies. Following data collection, highly accurate (Q43, >99.99%) circular consensus sequences (“HiFi reads”) were generated on the system. These HiFi data were used for all downstream analyses. HiFi read sequences were de-multiplexed by known barcode sequences and mutational analyses performed. To generate the consensus sequence of the seeding population, RNA isolated from a culture infected with 100 pfu of virus was used.

### Single Infectious Unit sequencing using nanopore sequencing

RNA was isolated from each well of infected cells cultured in a 96-well plate. Following cDNA synthesis, a 5.9 kb amplicon, encoding the nsP2-nsP3-nsP4 regions, was generated by PCR for each sample using the 9nt-barcoded nsP2-forward and nsP4-reverse primers described above (See Supplementary data, Sequences of the barcoded_primers.xlsx). Following amplicon generation, 2 µL from each PCR reaction were pooled together, purified with KAPA Pure magnetic beads (Roche) at a 0.5 × bead-to-sample volume ratio, and eluted in nuclease-free water. From the purified PCR sample, 1 µg was used as input into the Nanopore ligation sequencing library preparation kit for amplicons (Oxford Nanopore Technologies). If more than one 96-well plate worth of samples was to be sequenced together then the library preparation was done in conjunction with Nanopore native barcoding of each 96-well plate sample pool (Oxford Nanopore Technologies). Each sequencing run was performed using a MinION Mk1c sequencer with R9.4.1 flow cell for 6 hours with high accuracy basecalling following the sequencing run. After obtaining the raw fastq files, read quality check was performed using Fastqc. Read length distribution was examined to confirm the majority of reads fully cover the 5.9 kb amplicon. Sequencing data were de-multiplexed depending on the barcode sequences used for samples (see *Supportive information*, Sequences of the barcoded_primers.xlsx) with an in-house script using 9-nt long barcode sequence plus 10-nt long primer sequence for the de-multiplexing process.

### Sequence analysis with bioinformatics

Each demultiplexed sample fastq file was aligned to the reference genome sequence (VEEV GenBank: L01443.1) using Minimap2(Li, 2018), which is efficient for aligning long sequencing reads. To identify mutated sequences, we used mpileup from samtools (now bcftools)(Li et al., 2009). From each mapped bam file, the reference and alternative nucleotide sequences for each position of the target region were counted and saved in VCF format. The alternate allele frequency is used as the mutation frequency

The significance of the identified mutations was assessed with the Fisher’s exact test to calculate the p-value and Benjamini–Hochberg procedure to get false discovery rate (FDR). Occurrences of reference sequences and alternate sequences at each position of the target region from the control samples were counted to set these values as the baseline sequencing error frequency. Similarly, these occurrences of reference and alternate sequences from the treated samples were counted to set the observed allele frequency at each position. Using these values, we constructed a 2 × 2 table to compute p-value per each position. The FDR values were complied with the Benjamini-Hochberg procedure with the p-values of all loci (Benjamini & Hochberg, 1995). The overall mutation detection pipeline is depicted in *Supportive Method Figure 1*.

### Single infection unit viral growth kinetic assay

Vero 76 cells were seeded in an imaging culture media (1X MEM-E with 2 % FBS and 25 mM of HEPES without phenol red) in a 384-well plate at 5,000 cells/well and cultured overnight in a CO_2_ incubator. The cell plate was infected with virus at approximately 0.5 TCID50/well and incubated at 37 °C in the Cytation 5 (BioTek) supplemented with 5% CO_2_. The amount of virus per well was validated by a back titration of the diluted samples. The expression of EGFP was monitored every 20 min for 72 hrs. The growth analysis was performed using the fit function of AMiGA program(Midani et al., 2021) with additional arguments of ‘ --interval 1200 --do-not-log-transform -sfn 30’. Kinetic parameters were calculated with EGFP readouts (RFU) and the data were normalized based on the control group with the normalize function of AMiGA with an argument of --normalize-by. Wells at the two outermost positions (i.e., columns of 1,2,23, and 24, and rows of A, B, O, and P) were omitted for analysis for potential edge effects. Wells with the maximum growth rate (gr) > 200/hr and k-error threshold < 20% were defined as positive in viral replication and selected for further analyses.

### Selection and determination of ML336-resistant clones

For a bulk testing, Vero76 cells were seeded in 6-well plates at a density of 500,000 cells/well and cultured overnight. The next day, cells were infected with 50,000 pfu/well of virus for one hour at 37 °C and then rinsed with 2 mL of PBS. Then cells were overlayed with an agarose overlay media (0.6% agarose in MEM-E with 10% FBS), containing 1 µM of ML336 and cultured in a 37 °C with 5% CO_2_ incubator for two days. After cells were fixed with 4% paraformaldehyde, agar plugs were removed, and viral plaques were visualized with a counter staining with 0.05% neutral red solution. For individual isolation and selection, Vero 76 cells seeded in 384-well plates were infected with TC-83 virus that were diluted to at 0.01 resistant mutant/well (100, 5, and 5 pfu/well for virus produced in 0, 1, and 10 µM-NHC conditioned virus).

### Viral growth kinetic assay of mutant virus

Vero 76 cells were seeded in a virus culture media (1X MEM-E with 2 % FBS and 25 mM of HEPES) in a 96-well plate at 24,000 cells/well and cultured overnight in a CO_2_ incubator. The cell plate was infected with virus at MOI of 10 or 0.1 (n=8) in a volume of 25 µL. After a 1 hour incubation at 37 °C, 75 µL of culture media was added for each well and the expression of EGFP was monitored every 20 min for 28 hours at 37 °C in Cytation 5 (BioTek) provided with 5% CO_2_. The growth analysis was performed as described above with the AMiGA program(Midani et al., 2021).

## ACKNOWLEDGEMENT

We thank Firas Midani for helping with the growth analysis using AMiGA. The present work has benefited from University of Louisville Sequencing Technology Center.

## SUPPORTIVE INFORMATION

**Supportive Figure 1:**
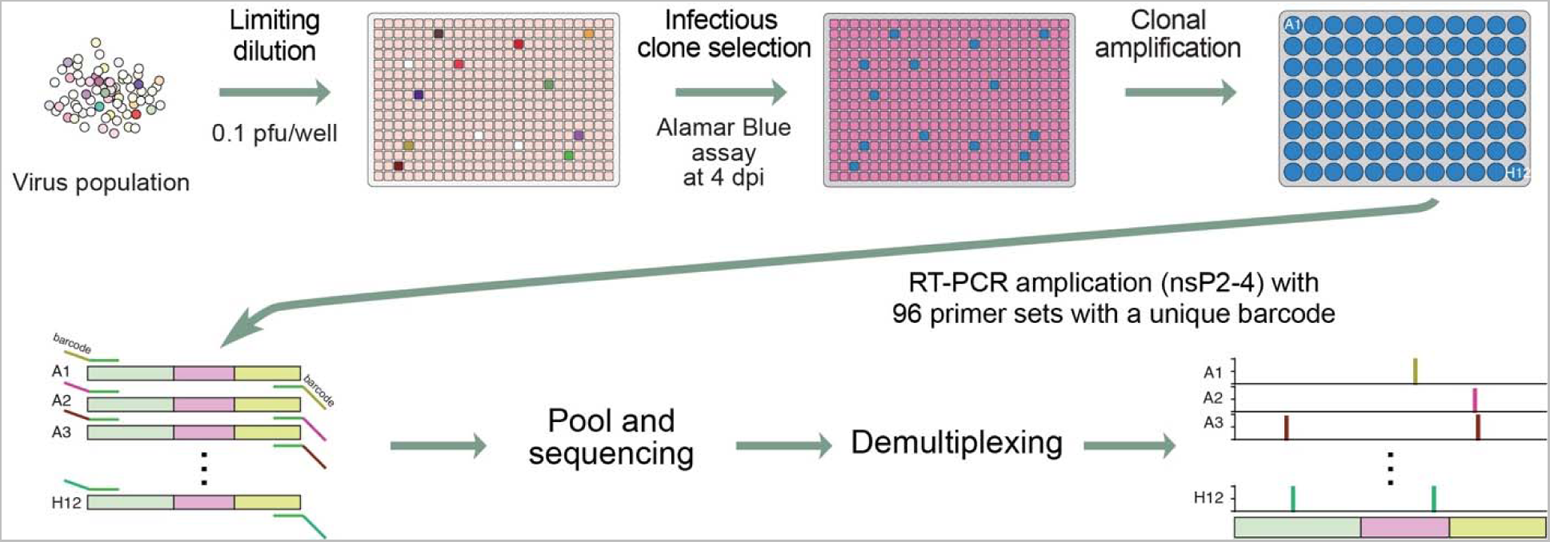
The overall mutation detection bioinformatics pipeline

**Supportive Figure 2:**
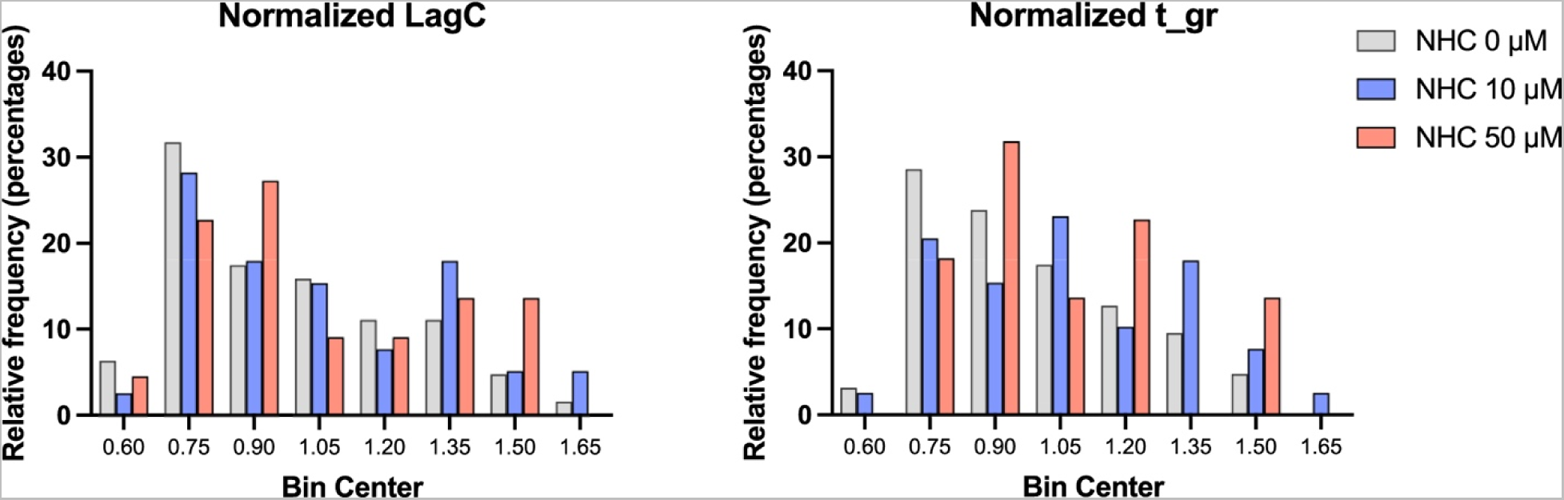
Analysis of kinetic parameters of virus clones conditioned with NHC.

**Supportive Method Figure 1:**
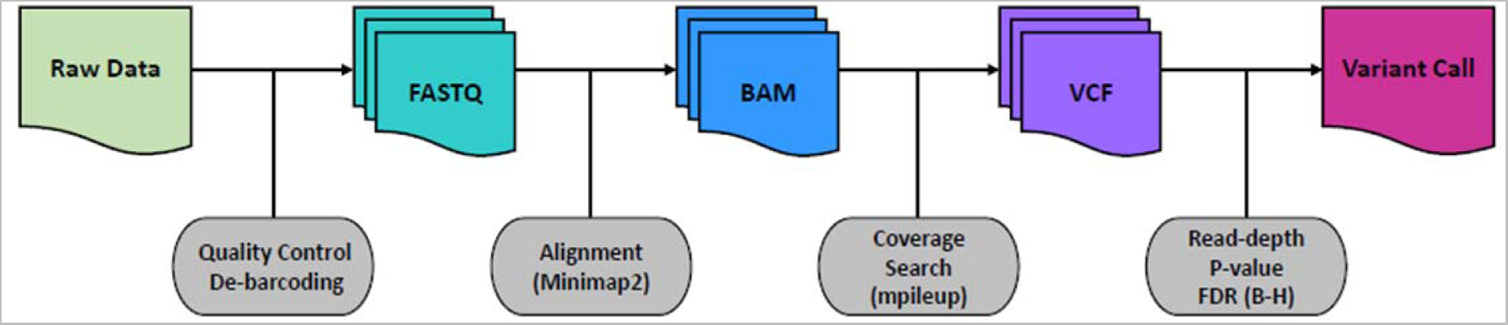
The overall mutation detection bioinformatics pipeline

**Supportive Table 1.**
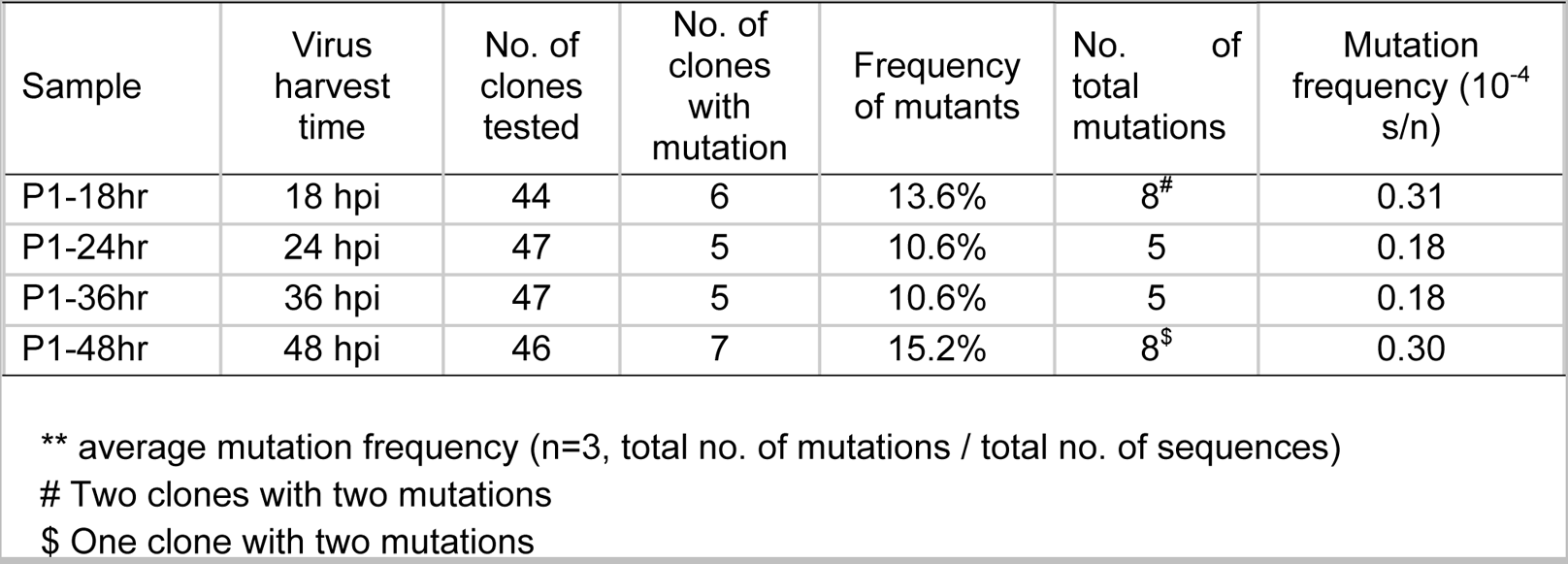
Mutation frequency of normal VEEV population

**Supportive Table 2.**
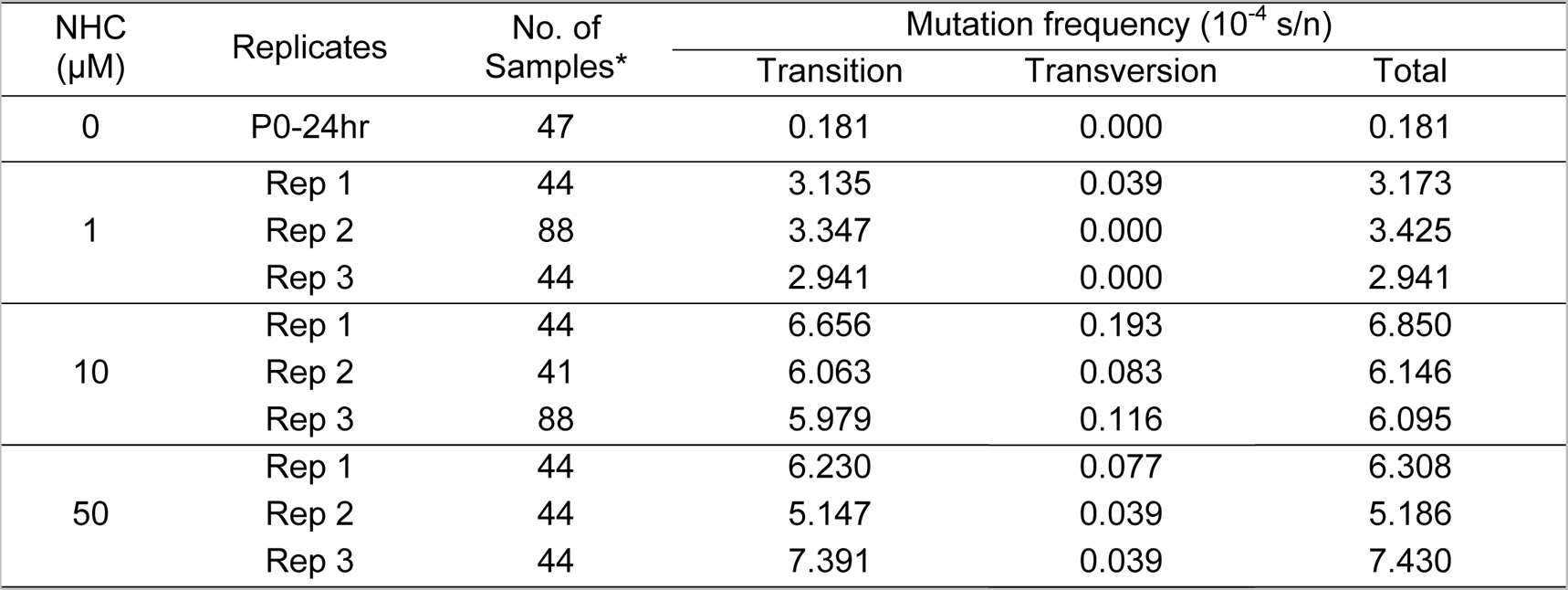
Analysis of mutation in response to NHC treatment.

**Supportive Method Table 1:**
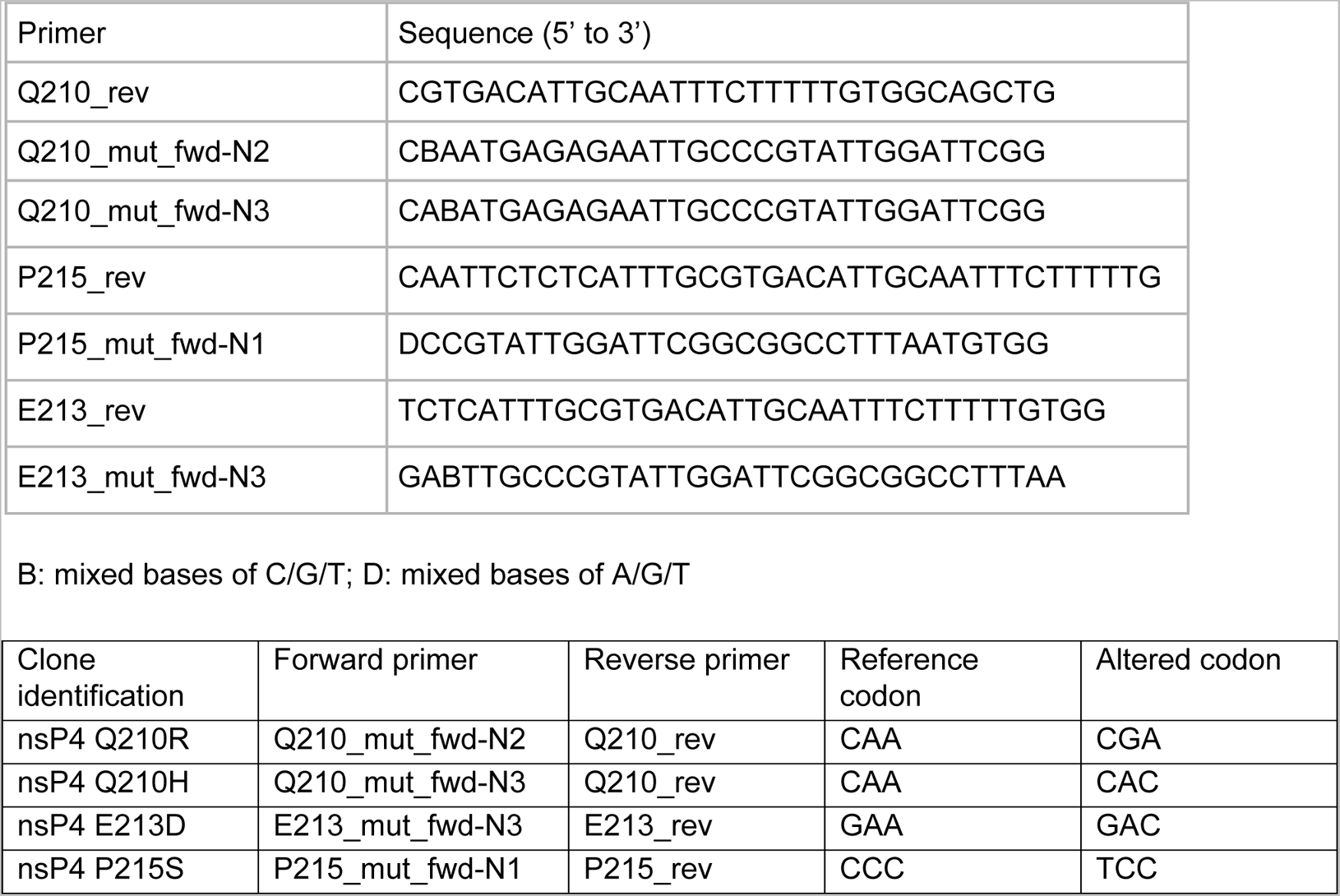
Primers and Their Sequences Used For a Site-Directed Mutagenesis

